# Spatial localization of CD16a at the human NK cell ADCC lytic synapse

**DOI:** 10.1101/2024.08.09.605851

**Authors:** Patrick Ross, Hijab Fatima, Dan P. Leaman, Jessica Matthias, Kathryn Spencer, Michael B. Zwick, Scott C. Henderson, Emily M. Mace, Charles Daniel Murin

**Affiliations:** San Diego Biomedical Research Institute, San Diego, CA, USA; Department of Integrative Structural and Computational Biology, Scripps Research, La Jolla, CA, USA; Department of Pediatrics, Columbia University Irving Medical Center, New York, NY, USA; Department of Immunology and Microbiology, Scripps Research, La Jolla, CA, USA; Abberior Instruments America, Bethesda, MD, USA; Core Microscopy Facility, Scripps Research, La Jolla, CA, USA; Department of Molecular Medicine, Scripps Research, La Jolla, CA, USA

## Abstract

Natural Killer (NK) cells utilize effector functions, including antibody-dependent cellular cytotoxicity (ADCC), for the clearance of viral infection and cellular malignancies. NK cell ADCC is mediated by Fc*γ*RIIIa (CD16a) binding to the fragment crystallizable (Fc) region of immunoglobulin G (IgG) within immune complexes on a target cell surface. While antibody-induced clustering of CD16a is thought to drive ADCC, the molecular basis for this activity has not been fully described. Here we use MINFLUX nanoscopy to map the spatial distribution of stoichiometrically labeled CD16a across the NK cell membrane, revealing the presence of pairs of CD16a molecules with intra-doublet distance of approximately 17 nm. NK cells activated on supported lipid bilayers by Trastuzumab results in an increase of synaptic regions with greater CD16a density. Our results provide the highest spatial resolution yet described for CD16a imaging, offering new insight into how CD16a organization within the immune synapse could influence ADCC activity. MINFLUX holds great promise to further unravel the molecular details driving CD16a-based activation of NK cells.

## Introduction

Natural Killer (NK) cells are innate immune lymphocytes that play important roles in the clearance of viral infection and cellular malignancies.^1^ A critical effector function of NK cells is antibody-dependent cellular cytotoxicity (ADCC) in which Fc*γ*RIIIa (CD16a) binds to the fragment crystallizable (Fc) domain of immunoglobulin G (IgG) within immune complexes on the target cell surface, inducing NK cell activation and degranulation.^2^ ADCC activity is central to the efficacy of many therapeutic monoclonal antibodies (mAbs),^3–5^ which can be optimized for enhanced ADCC activity. Despite the importance of spatial modulation for activating receptor function, a detailed molecular description of CD16a spatial distribution within the cellular context has not been described.

CD16a is a type I, single pass transmembrane glycoprotein with an intracellular domain and an extracellular domain containing membrane proximal and distal Ig-like domains. CD16a does not contain any signaling motifs in the intracellular domain, but instead signals through association with homo- or hetero-dimers of CD3ζ^6,7^ and/or FcR*γ*,^8,9^ which contain an immunoreceptor tyrosine-based activation motif (ITAM) on their intracellular tails.^10^ Glycosylation of both the extracellular domain of CD16a and IgG Fc influences binding affinity, consequently altering CD16a activity.^11^ Mechanistically, it is hypothesized that CD16a clustering is necessary for immune cell activation.^12^ While many activating receptors utilize ligand induced oligomerization to stimulate downstream signaling, the molecular details surrounding CD16a-based activation are incompletely understood. A recent study investigating murine bone marrow derived macrophages suggested that CD16a dimerization could induce inhibitory signaling through inhibitory ITAM signaling,^13^ proposing that a dimer may be the resting or inhibitory state.

NK cell cytolytic activity occurs through cytoskeletal rearrangements that result in the formation of an immunological synapse between the NK cell and target cell.^14^ At the immunological synapse, NK cell receptor rearrangement induces receptor signaling and NK cell activation.^15^ Optical microscopy has provided key insights into the dynamics and protein interactions occurring at the NK cell immunological synapse. Advancements in light microscopy now allow for the localization of single fluorophores attached to biomolecules with molecular precision by probing the fluorophore position with a focal excitation pattern containing an intensity minimum. MINFLUX localization usually uses the zero intensity of a donut shaped excitation beam to gradually approach the fluorophore position. The decreasing fluorophore-to-zero distance enhances the positional information content of each detected photon, rendering MINFLUX the most photon-efficient super-resolution microscopy concept to date based upon the detection of a minimal photon flux.^16–19^ Application of MINFLUX has facilitated the ability to better understand protein dynamics such as the walking of kinesin along microtubules^20–22^ and the conformational changes of PIEZO-1.^23^ These examples highlight the potential of MINFLUX to illuminate new structural and functional biology previously unattainable with alternative super-resolution techniques.

Here we leverage the precise molecular localization of MINFLUX nanoscopy to describe the spatial distribution of CD16a on the NK cell membrane. Using a genetically introduced CD16a-SNAP construct, we observed pairs of CD16a molecules across the NK cell membrane with a spacing between fluorophores of about 17 nm.

## Results

### Generation and functional validation of an NK cell line with tagged CD16a

To investigate the spatial organization of CD16a in the NK cell membrane during the process of ADCC, we generated an NK-92 cell line with a genetically integrated CD16a gene containing a SNAP-tag on the C-terminus (Fig. 1A and Fig. S1) and a control cell line with an untagged version of CD16a. NK-92 cells do not express CD16a, ensuring that our construct would be the only CD16a expressed on the cell surface. The construct was introduced into the NK-92 genome via nucleofection with a piggyBac transposase.^24–27^ After recovery from transfection, the cells were sorted by FACS and expanded, resulting in an NK-92 cell line expressing the CD16a constructs, which we termed NK-92^CD16^ and NK-92^CD16-SNAP^ (Fig. 1B).

**Figure 1:**
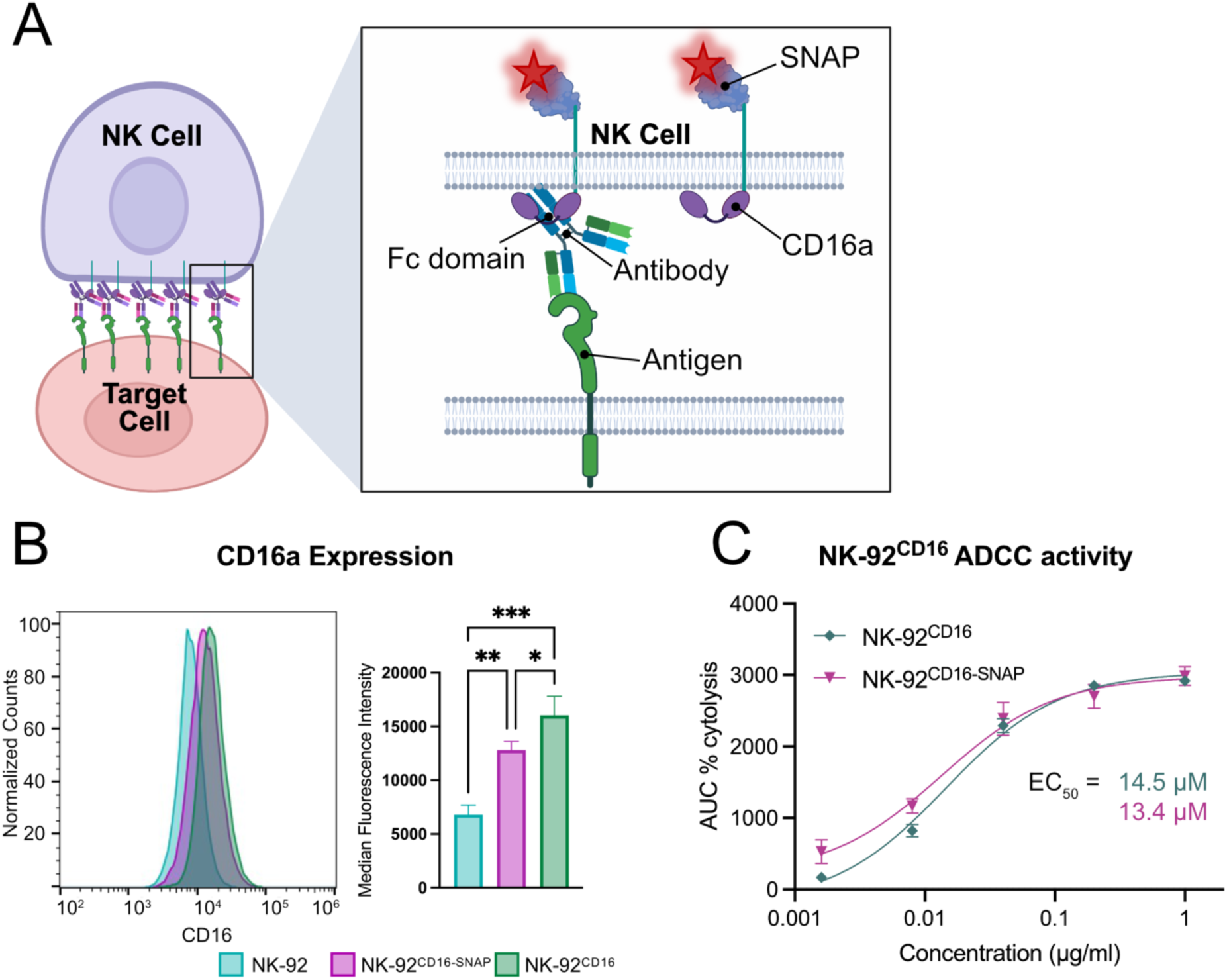
NK-92^CD16-SNAP^ does not disrupt ADCC activity. (A) Cartoon representation of the NK cell immunological synapse formed with an antibody and representation of the SNAP-tag used for cell labeling of CD16a with AF647. (B) Flow cytometry quantification of CD16a expression on NK-92, NK-92^CD16-SNAP^, and NK-92^CD16^ cells stained with α-CD16 antibody 3G8 (AF488 labeled). (C) Trastuzumab dose response curves of ADCC activity for NK-92^CD16-SNAP^ and NK-92^CD16^ positive NK cells. AUC = area under the curve. *** = *p* < 0.001; ** = p < 0.01; * = p < 0.05.

To validate ADCC activity of the newly created cell lines, we used an impedance-based assay that provides both a measurement of the rate of killing and maximum cytolytic activity through changes in cell morphology of adherent target cells.^28^ We used SKOV-3 cells as targets, which express HER2 and can be targeted for ADCC via Trastuzumab.^29^ We observed similar cytolytic activity whether NK-92 cells expressed CD16a alone or the CD16-SNAP construct, and both cell lines displayed comparable activity to NK cells isolated from human peripheral blood mononuclear cells (PBMCs) (Fig. 1C and Fig. S2).^30^

### Antibody cross-linking induces phosphorylation of CD3ζ in NK-92^CD16-SNAP^ cells

To establish the utility of the NK-92^CD16-SNAP^ cell line for downstream microscopy studies, we modeled the NK cell immune synapse using antibody coated glass to achieve receptor cross-linking, similar to previous studies.^31,32^ For this experiment, we compared synapses formed on glass coated with both anti-CD16 (α-CD16) and α-CD18 (anti-lymphocyte function-associated antigen-1, or LFA-1) antibodies to induce cell spreading (Fig. 2A), with an α-CD18 antibody alone (Fig. 2B), or with poly-L-lysine (PLL) as a negative control (Fig. 2C). NK-92^CD16-SNAP^ cells were fixed and stained with a SNAP ligand, which labels SNAP-tagged CD16a, and an antibody targeting phosphorylated CD3ζ (pCD3ζ) as a marker of downstream activation^33–35^ that can be imaged by confocal microscopy (Fig. 2A-C).

**Figure 2:**
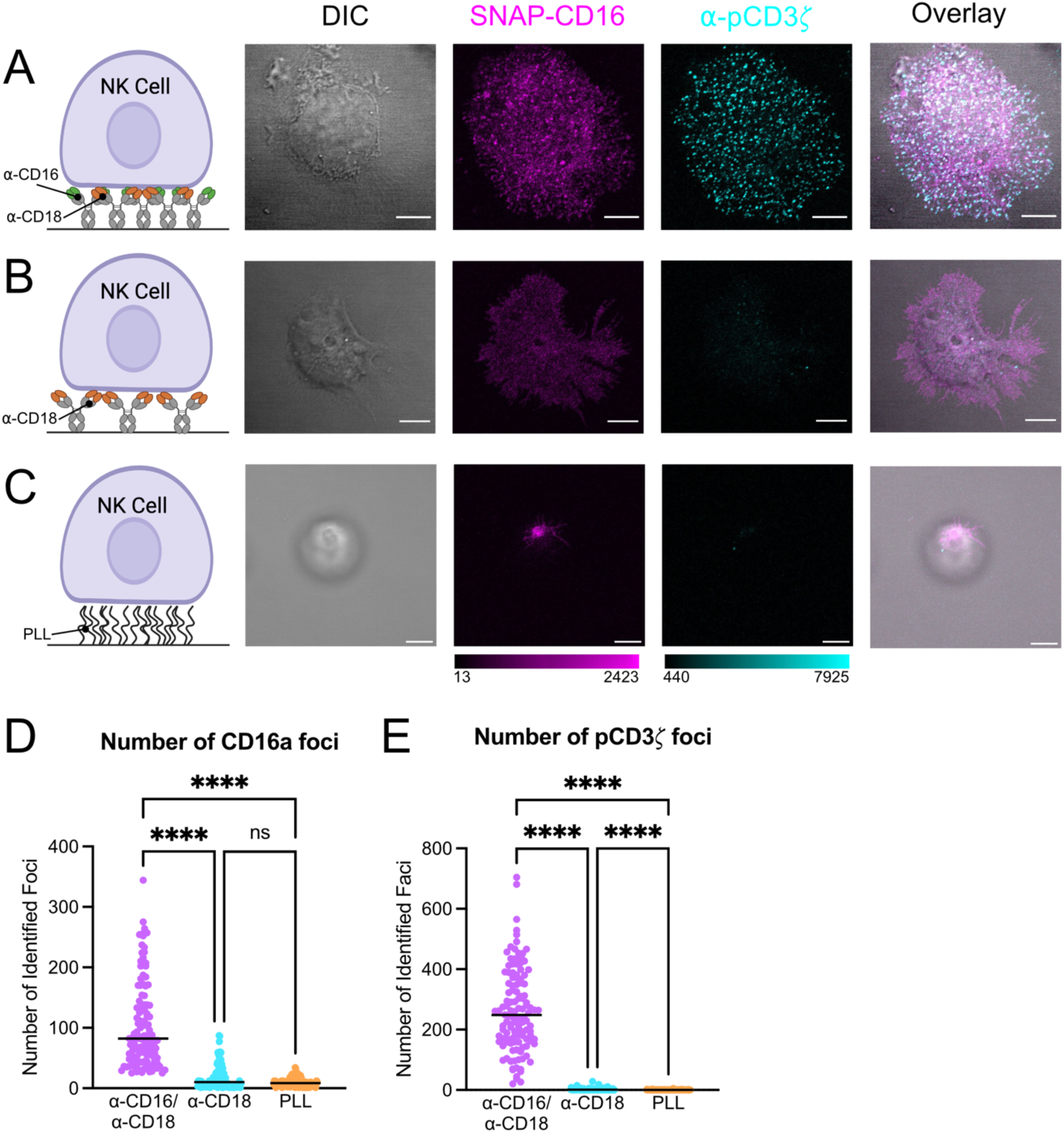
Antibody cross-linking of CD16a and CD18 on glass coverslips induces phosphorylation of CD3ζ. (A-C) The far-left column depicts a cartoon representation of activating conditions used to model CD16a mediated activation. Then from left to right shows representative differential interference contrast and confocal micrographs of SNAP-CD16 and pCD3ζ on NK-92^CD16-SNAP^ synapses formed on glass coated with (A) α-CD16 and α-CD18 antibodies, (B) α-CD18 antibody, or (C) PLL. (D) Presents quantification of the number of pCD3ζ foci per cell. (E) Displays quantification of the number of CD16a foci identified. Scale bars represent 5 µm. **** = *p* < 0.0001, ns = not significant.

The NK^CD16-SNAP^ cells ligated on glass dual-coated with α-CD16/α-CD18 antibodies produced clearly discernable foci of pCD3ζ where the artificial synapse had been formed (Fig. 2A). The number of CD16a foci in NK cells was significantly higher on glass coated with α-CD16/α-CD18 antibodies compared to the α-CD18 antibody solo-coating or the negative control (Fig. 2A-C and Fig. 2D). Additionally, we observed a greater number of pCD3ζ foci in NK cells activated on glass with α-CD16/α-CD18 antibodies than with CD18 cross-linked or resting (PLL) NK cells (Fig. 2A-C and 2E). This reflects similar observations for cross-linking CD16a on supported lipid bilayers.^31,36^ Taken together, these data demonstrate that NK cells activated through cross-linking of CD16a and CD18 results in the formation of regions with dense molecular clusters of CD16a. Conversely, on NK cells activated through CD18 cross-linking and resting NK cells, CD16a is more diffusely spread across the cell membrane.

### MINFLUX nanoscopy reveals pairs of CD16a

To define how CD16a is distributed across the NK cell membrane with molecular precision, we turned to MINFLUX nanoscopy. For our initial investigation, we used the same conditions that we had validated by confocal microscopy in the first step (see above). NK-92^CD16-SNAP^ cells were activated through cross-linking of either α-CD16a/α-CD18, α-CD18 alone, or incubated on PLL (Fig. 3A-C and Fig. S4). Our MINFLUX analysis pipeline (Fig. S3) mapped individual CD16a molecules as groups of several localizations and throughout our data collection, we regularly obtained a mean localization precision of ∼2.5 nm (Fig. S4). To determine if CD16a forms clusters, we used Ripley’s H in which a value >0 represents increased clustering and a value <0 represents dispersion. From this data, we observed in all the conditions that CD16a is not randomly distributed across the cell membrane, but in clusters (Fig. 3D).^37^ On average, we saw little variation in the degree of randomness between the experimental conditions; however, in some regions of interest (ROIs) in which MINFLUX data was acquired, we observed greater clustering compared to other regions (Fig. 3D), indicating heterogeneity in CD16a distribution across different cells.

**Figure 3:**
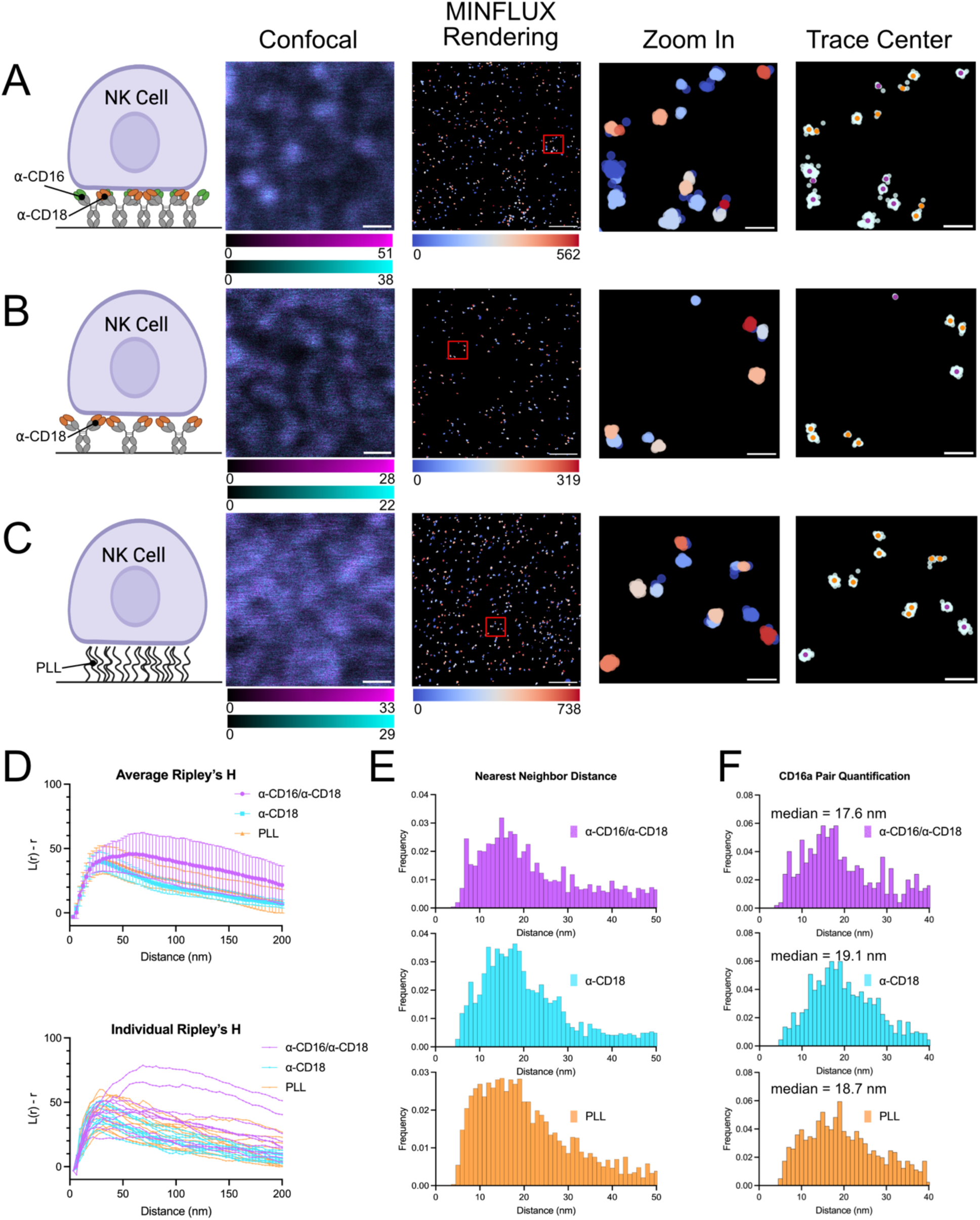
CD16a in an artificial immune synapse on antibody coated glass forms pairs. (A-C) The far-left column depicts a cartoon representation of activating conditions used to model CD16a mediated activation and displays representative images of NK-92^CD16a-SNAP^ confocal and MINFLUX data of immune synapses on coated glass. From left to right, images show confocal reference, MINFLUX data colored given through DBSCAN, a zoomed in region of MINFLUX localizations as marked in the overview, and trace centers (single localizations are colored in purple, and isolated pairs are in orange) overlayed on raw MINFLUX data (colored in light cyan). Conditions are glass coated with (A) α-CD16 and α-CD18 antibodies, (B) α-CD18 antibody, and (C) PLL. (D) Average Ripley’s H function (top) and individual region of interest (ROI) Ripley’s H function (bottom) of MINFLUX data. (E) Nearest neighbor analysis of all MINFLUX data. (F) Nearest neighbor analysis of isolated pairs of CD16a molecules. (G) The total number of isolated CD16a pairs observed in all ROIs. Scale bars in confocal images and raw MINFLUX renderings represent 500 nm and zoomed in region scale bars represent 50 nm.

Strikingly, MINFLUX imaging consistently visualized pairs of CD16a molecules across all ROIs collected for the three sample conditions. Looking more closely at the intermolecular CD16a distance, we observed that on average CD16a molecules of cells on glass coated with α-CD18 antibodies or PLL resided closer together than in presence of the α-CD16 antibody (Fig. 3E). We determined the nearest neighbor distance between isolated CD16a pairs (Fig. 3F) and observed a median nearest neighbor distance of 17.6 nm, 19.1 nm, and 18.7 nm of isolated pairs on NK cells activated on glass coated with α-CD16a/α-CD18, α-CD18, or incubated on PLL, respectively. Most notably, isolated CD16a pairs were more abundant in the CD18 cross-linking and PLL samples.

These initial results demonstrated the ability of MINFLUX to describe the molecular distribution of CD16a and identify potential new interactions not previously described for CD16a on NK cells. However, we questioned if activation of NK cells through CD16a cross-linking reflects the molecular nature of NK cell activation during ADCC, during which CD16a is engaged through ligand binding to an Fc domain in the context of an immune complex rather than cross-linked by an α-CD16 antibody. Further, the 3G8 clone used in the present study to activate NK cells is known to directly compete for binding with IgG.^38^ Therefore, we next sought to recapitulate a more realistic synaptic model to determine the molecular location of CD16a during NK cell ADCC.

### NK cells on supported lipid bilayers with mAb induces increased phosphorylation of CD3ζ

To design a model that would recapitulate the immune synapse during ADCC, we used a supported lipid bilayer (SLB) displaying both an antigen of interest and intercellular adhesion molecule 1 (ICAM-1), which is the ligand for LFA-1 on NK cells that promotes adhesion and synapse formation.^31,39^ We chose to develop our system using HER2 as the target antigen, which is therapeutically targeted by NK cell-mediated ADCC with Trastuzumab.^40^ We generated SLBs using 1-palmitoyl-2-oleoyl-glycero-3-phosphocholine (POPC) and (1,2-dioleoyl-sn-glycero-3-[(N-(5-amino-1-carboxypentyl)iminodiacetic acid)succinyl] (nickel salt)) (NTA(Ni)-containing) lipids. After addition of His-tagged HER2 and ICAM-1, bilayer fluidity was confirmed by fluorescence recovery after photobleaching (FRAP) (Fig. S5).

We next confirmed that our model of a target cell membrane would induce antibody-mediated NK cell activation. As a marker of NK cell activation, we again used an increase in pCD3ζ as measured by confocal microscopy. We incubated NK-92^CD16-SNAP^ cells on bilayers containing HER2 and ICAM-1 and compared them to NK-92^CD16-SNAP^ cells on bilayers that had been opsonized with Trastuzumab (Fig. 4A-B). We observed that the addition of Trastuzumab resulted in an increase in the number, size, and intensity of pCD3ζ foci as observed by confocal microscopy (Fig. 4C-E). This suggests that the SLB induced NK cell activation through CD16a specifically. Visually, we observed no difference in CD16a distribution between the two experimental conditions, which contrasted with our glass-activated data showing regions of enhanced CD16a fluorescence intensity on NK cells cross-linked via the α-CD16 antibody. Therefore, we further studied the underlying molecular distribution of CD16a on NK cells activated by immune complexes rather than activated via cross-linking by the α-CD16 antibody, and whether CD16a cross-linking/clustering is indeed required for antibody dependent cellular activation.

**Figure 4:**
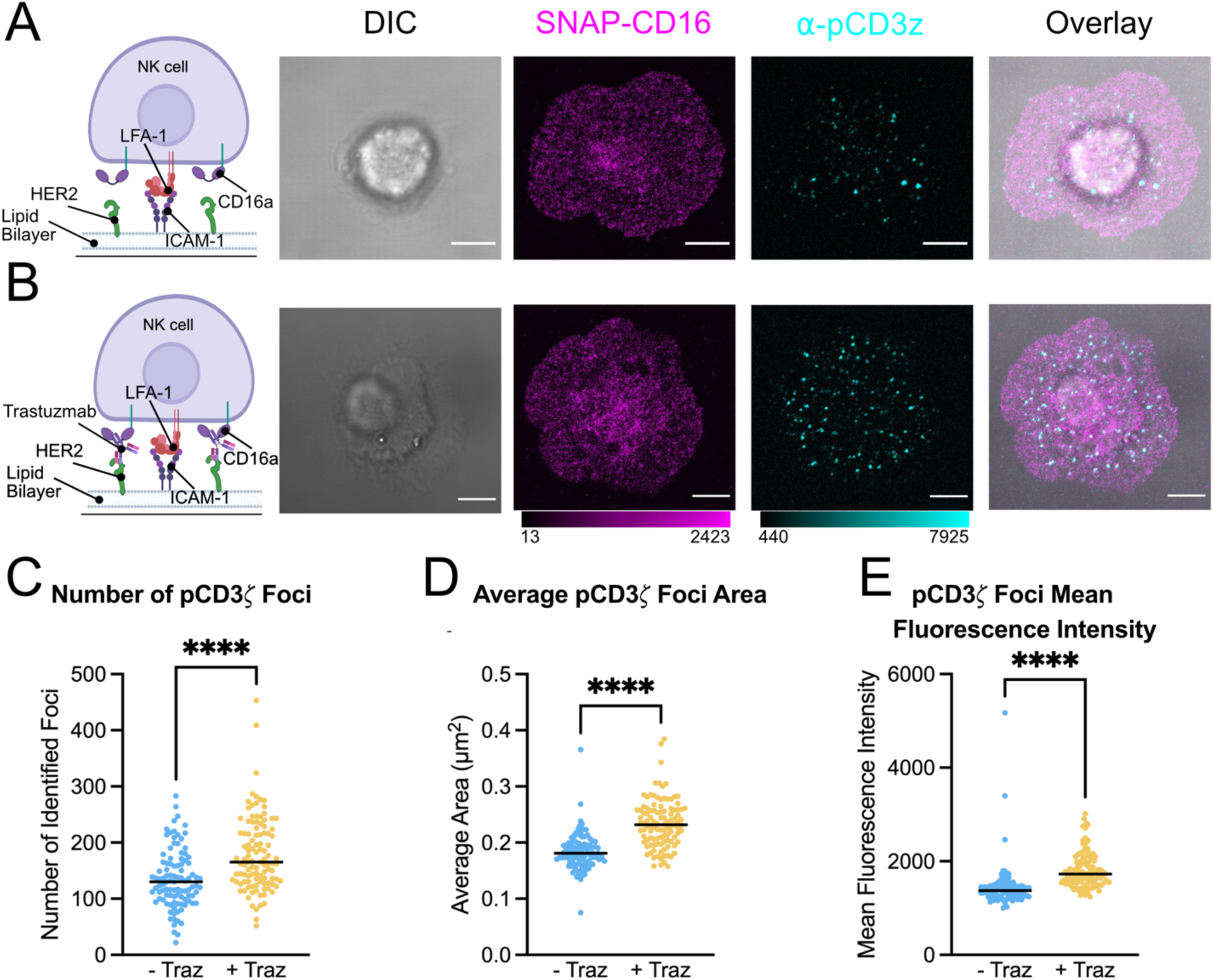
SLB displaying HER2 and ICAM-1 opsonized with Trastuzumab induces phosphorylation of CD3ζ. (A-B) The far-left column depicts a cartoon representation of activating conditions used to model CD16a mediated activation. Shown are representative differential interference contrast and confocal micrographs of SNAP-CD16 and pCD3ζ on NK-92^CD16-SNAP^ synapses formed on SLBs containing HER2 and ICAM-1 (A) without Trastuzumab (-Traz) and (B) with Trastuzumab (+ Traz). Quantification of the number (C), average area (D), and fluorescence intensity (E) of foci of pCD3ζ per cell. Scale bars represent 5 µm. **** = *p* < 0.0001.

### MINFLUX imaging reveals a slight change in CD16a distribution on NK cell synapses formed on a SLB

We subsequently visualized the molecular distribution of CD16a in the context of artificial immune synapses between NK-92^CD16-SNAP^ cells and SLBs containing ICAM-1 and HER2 with MINFLUX nanoscopy (Fig. 5A-B and Fig. S4). Both in the presence and absence of Trastuzumab, CD16a was not randomly distributed across the cell membrane in the regions imaged (Fig. 5C). Further, we observed little difference in the degree of clustering between the two conditions based on Ripley’s H function (Fig. 5C).

**Figure 5:**
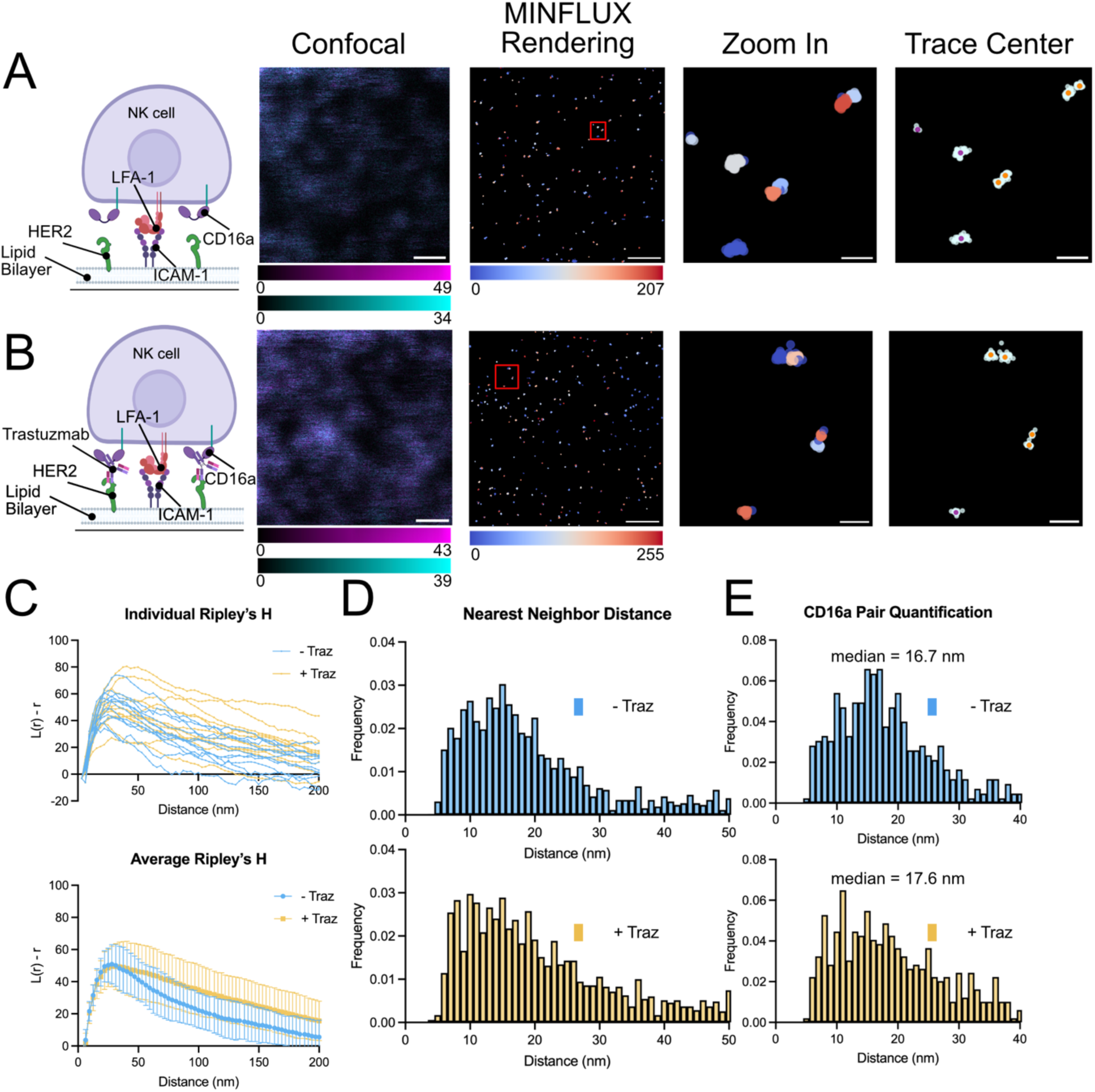
Slight change in CD16a distribution is observed in NK-92^SNAP-CD16^ artificial synapses on SLBs. (A-B) The left column depicts a cartoon representation of activating conditions used to model CD16a mediated activation. Representative images of NK-92^CD16-SNAP^ confocal and MINFLUX data of immune synapse on SLBs. From left to right, images show confocal reference, MINFLUX data colored by ID given through DBSCAN, a zoomed in region of MINFLUX localizations as marked in the overview, and trace centers (single localizations are colored in purple, and isolated pairs are in orange) overlayed on raw MINFLUX data (colored in light cyan). SLBs contained HER2 and ICAM-1 (A) without Trastuzumab (-Traz) and (B) with Trastuzumab (+ Traz). (C) Average Ripley’s H function (top) and individual ROI Ripley’s H function (bottom) of MINFLUX data. (D) Nearest neighbor analysis of all MINFLUX data. (E) Nearest neighbor analysis of isolated pairs of CD16a molecules. Scale bars in confocal images and raw MINFLUX data represent 500 nm and zoomed in region scale bars represent 50 nm. MINFLUX microscopy data were collected across at least three independent experiments with a total of 12 cells analyzed.

In all SLB data sets, we commonly observed the localization of CD16a occurred in pairs (Fig. 5A-B). Calculating nearest neighbor distances, we found little difference in the average nearest neighbor distance between the two conditions (Fig. 5D). When we isolated these pairs and performed nearest neighbor analysis, we again observed a nearest neighbor intra-fluorophore distance of 16.7 nm and 17.6 nm on NK cells activated without and with Trastuzumab, respectively (Fig. 5E), like our results obtained with NK cells synapses formed on glass (Fig. 3F). We did not observe dense clusters of CD16a as previously reported for other NK cell receptors,^41–43^ though when NK-92 cells were activated with Trastuzumab, we did observe an increase in the number of regions with greater than four localized CD16a molecules in a 40 nm radius by Density-Based Spatial Clustering of Application with Noise (DBSCAN)^44^ (Fig. S6).

## Discussion

Here we leverage MINFLUX nanoscopy to describe the single molecule arrangement of CD16a within resting and activated NK-92 cells modified to genetically express SNAP-tagged CD16a. These data demonstrate that CD16a is found in pairs on resting cells with tags separated by around 17 nm. The relative frequency of this organization was reduced on NK-92 cells on glass coated with α-CD16/α-CD18. Pairs of CD16a were also observed on NK cells resting on SLBs and in the presence of an ADCC-activating monoclonal antibody. Taken together, our single molecule observations suggest that pairs of CD16a molecules could represent a resting or inhibitory conformation that remains present even during NK cell activation. It is not clear from our current data if the pairs present in active NK cell synapses are also liganded to immune complexes, or if they are a subset of receptors that are not actively engaged by antigen. We should note that in these studies, we utilized SNAP-tagged CD16a to provide stoichiometric labeling and to help minimize the distance between the fluorophore and protein of interest. However, inherent labeling error provides a degree of uncertainty in precisely measuring the distance between receptors. Additionally, our ability to covalently conjugate and detect a fluorophore on every receptor could limit a complete description of CD16a spatial distribution.

The observed median inter-fluorophore distance of about 17 nm in CD16a pairs suggests that CD16a likely does not persist on the cell membrane as a homodimer but may instead be scaffolded by other proteins, such as CD3ζ (that is required for cell surface trafficking of CD16),^7,8^ that further separates the individual CD16a proteins (Fig. 6). It is possible that CD16a binding to an immune complex may induce a very subtle change in CD16a conformation that we cannot detect here, which could facilitate enhanced phosphorylation of CD3ζ and/or spatial rearrangement. Additional studies using the tools developed here, including multi-channel MINFLUX nanoscopy and structural biology techniques like cryo-EM, could help further elucidate the protein orchestration and structure of CD16a in resting NK cells and how additional signaling proteins interplay to direct ADCC-specific activation as opposed to other forms of NK cell activation.

**Figure 6:**
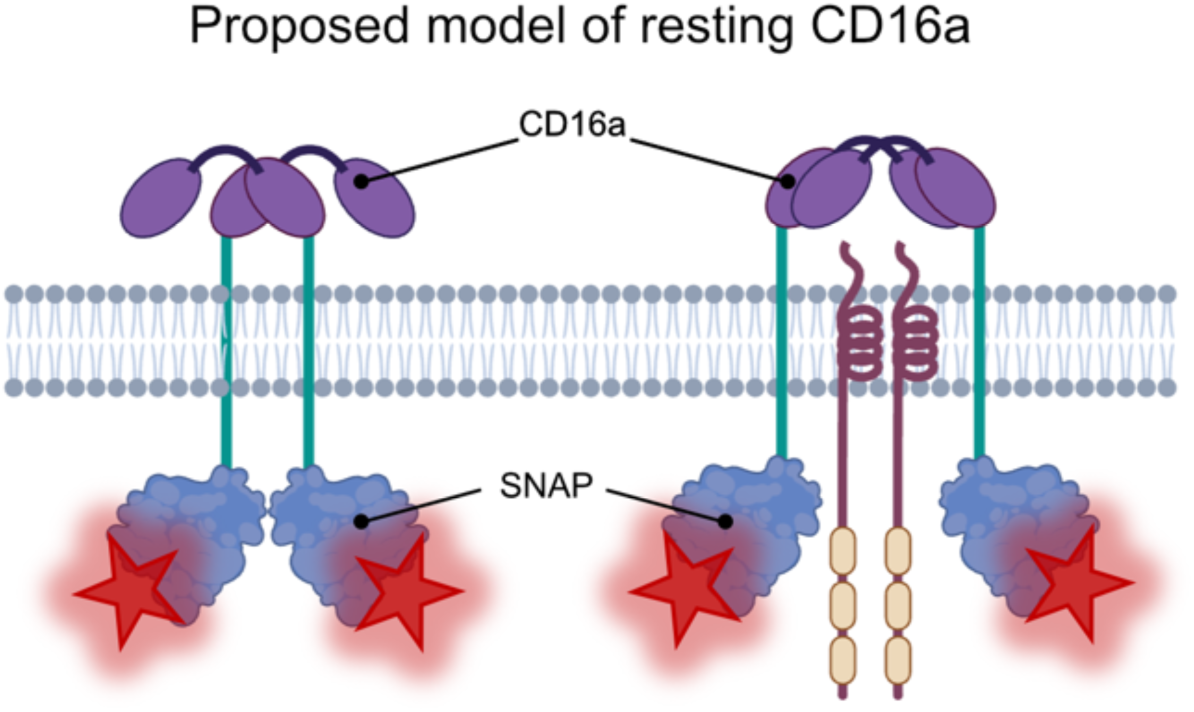
Working model of CD16a resting conformation. MINFLUX data of CD16a localization demonstrated that CD16a often is found in pairs with an intra-fluorophore distance of ∼17 nm apart. Shown is a cartoon representation of what this could look like in un-scaffolded CD16a (left) versus CD16a scaffolded by a homodimer such as pCD3ζ.

Our observation of CD16a pairs in all our samples sheds new light on the molecular mechanisms required for antibody-based activation via IgG receptors. Previous work suggests that activation of NK cells through CD16a is through the formation of CD16a clusters on the cell membrane.^31,36^ We did observe an increase in the frequency of areas with greater density of CD16a on activated NK cells, but the limited frequency of such regions restricts our current ability to describe their role in greater detail. While we are currently working to address this limitation in future models, these regions do not resemble the large, dense clusters observed for other NK cell receptors, such as KIR2DL1, KIR2DS1, or LILRB1.^41–43^ Our observations support a model where CD16a forms molecular clusters upon activation through slight changes in its distribution on the cell membrane, potentially influenced by other molecular interactions including other cell surface receptors or signaling partners (Fig. 6). Though not immediately clear, our observations could help to explain why there are disparities in antibody ADCC potency related to the target epitope.^45^

## Acknowledgements

All confocal and MINFLUX microscopy data were collected at the Scripps Research Microscopy Core facility. All cell sorting was performed at the Scripps Research Flow Cytometry Core facility. We would like to thank Dr. Otto Wirth, Dr. Clara Gürth and Abberior Instruments for assistance with MINFLUX acquisition, instrument and software troubleshooting, and data analysis. We would also like to thank Dr. Mark Patrick Farrell from the University of Kansas for providing the hyPBase plasmid; Dr. Andrew Ward at Scripps Research for the use of their equipment and feedback on this project and manuscript; and Dr. Dylan Owen for discussion on MINFLUX data analysis. Some figures were generated using BioRender.com through an academic subscription purchased by CDM.

## Author contributions

PR designed and performed all experiments, analyzed data, and wrote the manuscript. HF helped design confocal experiments and helped discuss and analyze microscopy data. DPL helped design SLB experiments and generated liposomes. JM helped design MINFLUX experiments and assisted with MINFLUX data collection and analysis. KS provided confocal and FRAP training, assisted with microscope operation, and provided confocal and FRAP data analysis training and assistance. MBZ provided SLB experimental design assistance. SCH provided MINFLUX training, microscopy experimental design, and assistance with MINFLUX data collection and data analysis. EMM provided assistance with microscopy experimental design and data analysis. CDM designed NK cell constructs, assisted with ADCC assays, helped with experimental design and data analysis, and helped to write the manuscript. All authors provided input toward editing the manuscript.

## Funding sum

Research reported in this publication was supported by the NIAID of the National Institutes of Health under award numbers 1R01AI167646 (CDM) and R01AI181784 (MBZ).

