## Supplemental Materials for "Spatial localization of CD16a at the human NK cell ADCC lytic synapse"

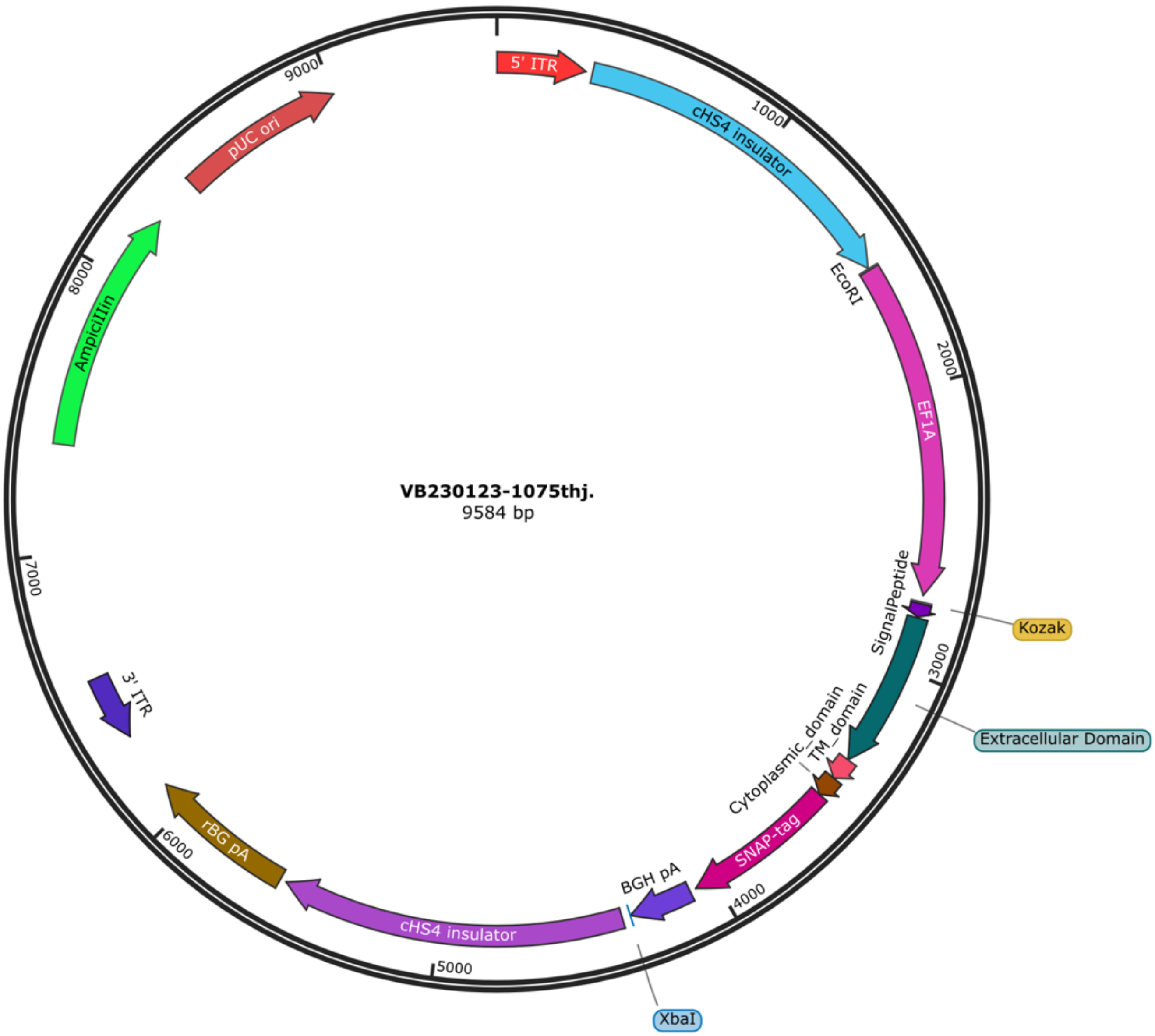


**Supplemental Figure 1: Plasmid map of CD16-SNAP plasmid.** Plasmid map used for generation of the NK-92$^{CD16\text{-}SNAP}$ cell line.

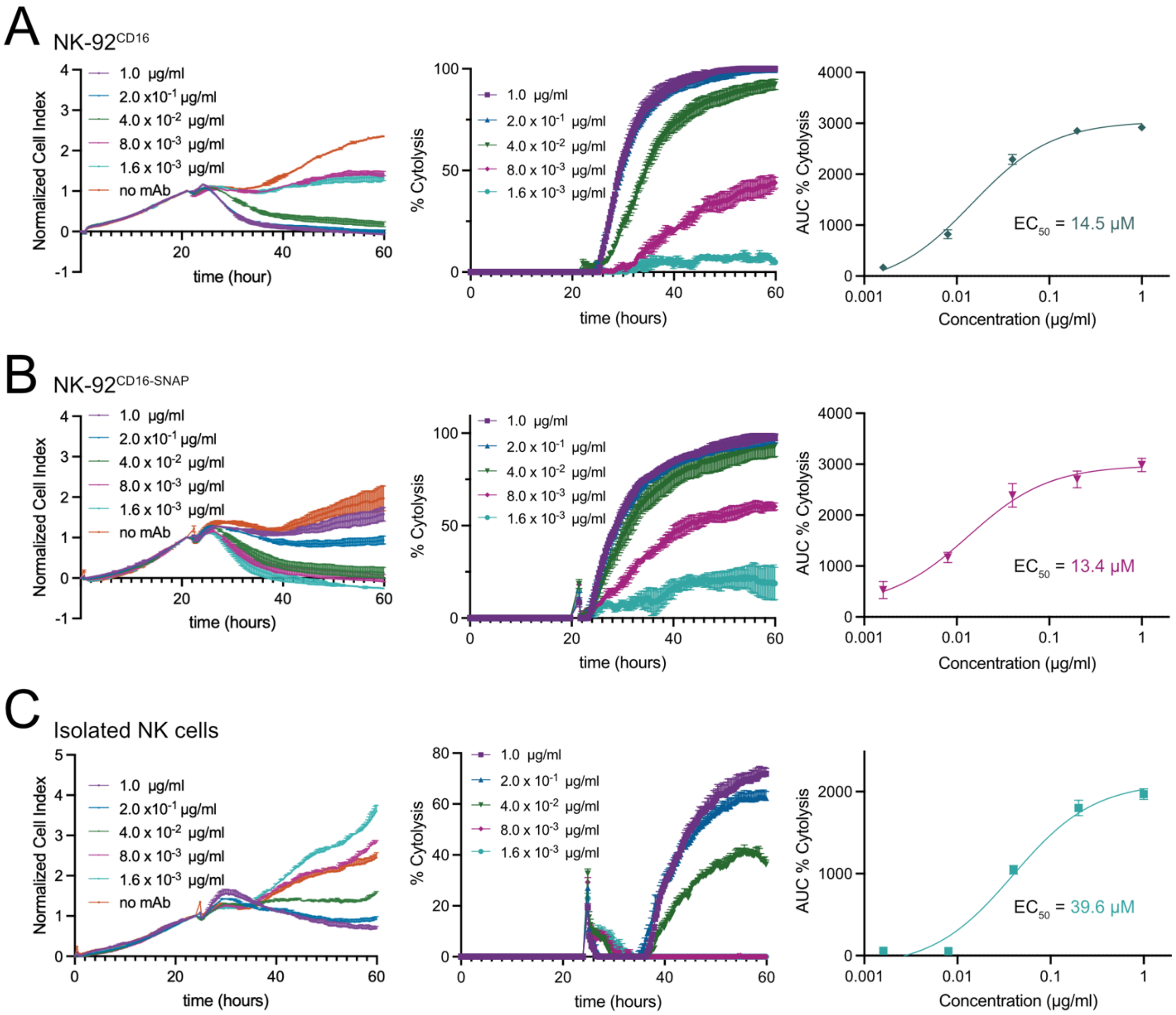


**Supplemental Figure 2: Comparison of NK-92$^{CD16}$, NK-92$^{CD16\text{-}SNAP}$, and isolated NK cell ADCC activity.** Representative data from impedance based ADCC activity. From left to right, raw normalized cell index, % cytolysis, and area under the curve (AUC) of % cytolysis for (A) NK-92$^{CD16}$, (B) NK-92$^{CD16\text{-}SNAP}$, and (C) isolated NK cells from PBMCs.

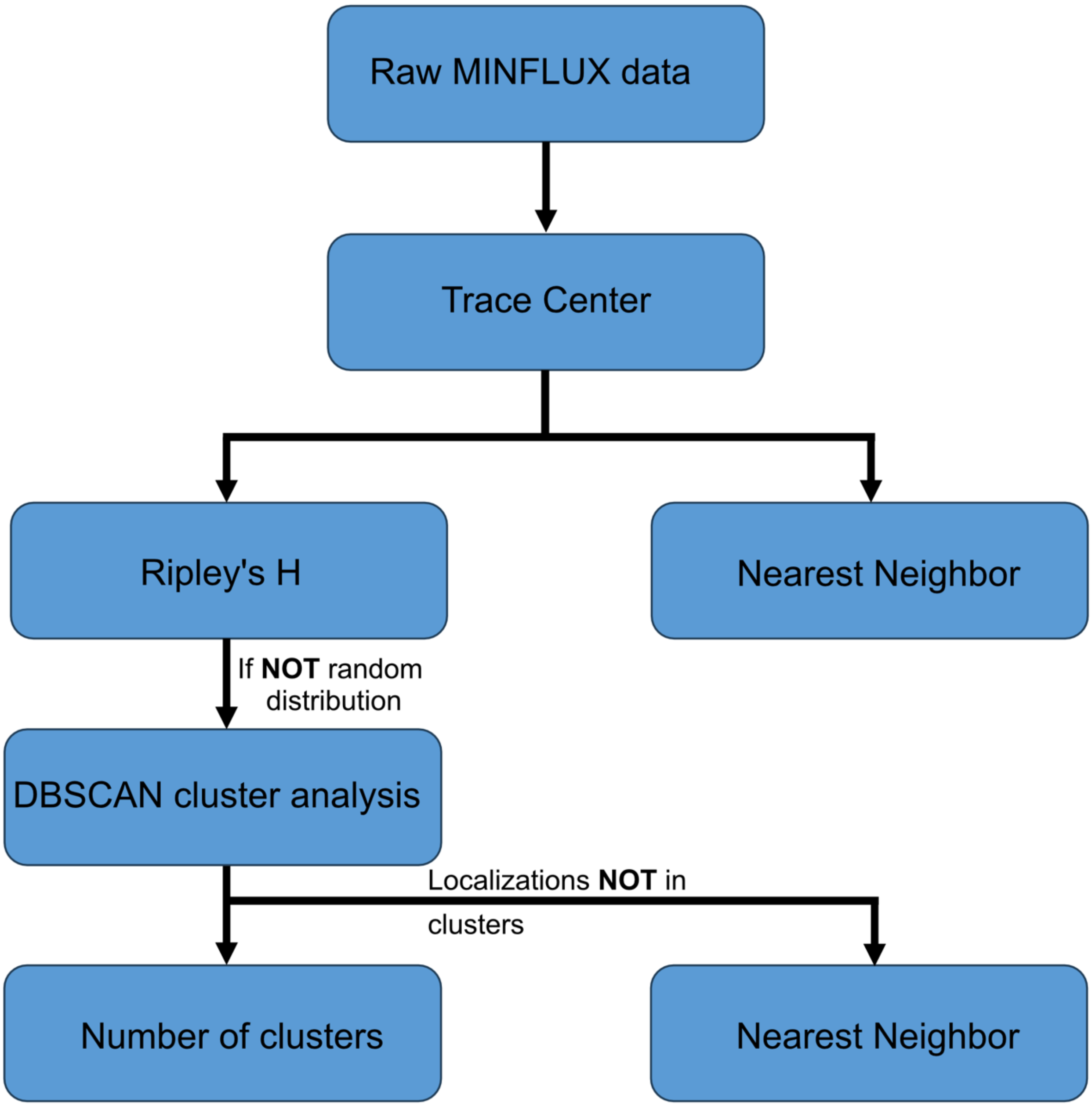


**Supplemental Figure 3: Data analysis workflow for MINFLUX nanoscopy collections.** Flow chart describing the data analysis pipeline used to analyze MINFLUX data. Raw MINFLUX data was first filtered, and the center of clusters identified provide the likely fluorophore localization. Fluorophore localizations were used to determine nearest neighbor distance. A second set of analysis of fluorophore localization was then used to determine if the data is clustered using Ripley's H. If the data was clustered, then clusters were identified using DBSCAN. The identified clusters were then counted. Secondary analysis was performed on localizations not in clusters, and pairs of localizations identified. Then the nearest neighbor distance between isolated pairs performed.

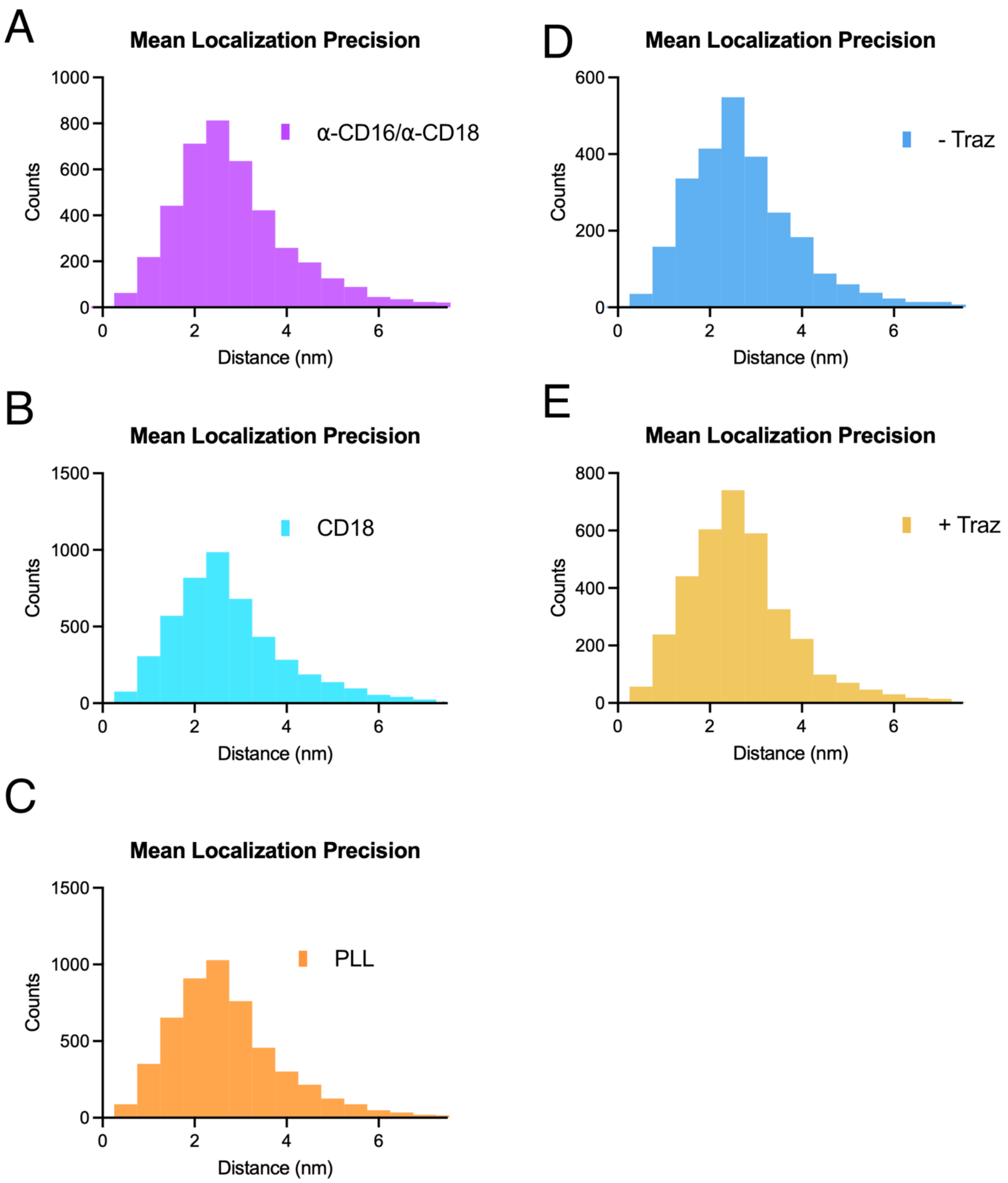


**Supplemental Figure 4: Mean localization precision of MINFLUX data.** (A) Representative mean localization precision of a data set acquired with cells on glass coated with α-CD16 and α-CD18 antibodies (4,146 trace centers). (B) Representative mean localization precision of a data set acquired with cells on glass coated with $\alpha$-CD18 antibody (4,744 trace centers). (C) Representative mean localization precision of a data set acquired with cells on glass coated with PLL (5,119 trace centers). (D) Representative mean localization precision of a data set acquired with cells on an SLB without antibody (2,564 trace centers). (E) Representative mean localization precision of a data set acquired with cells on an SLB opsonized with Trastuzumab (3,510 trace centers).

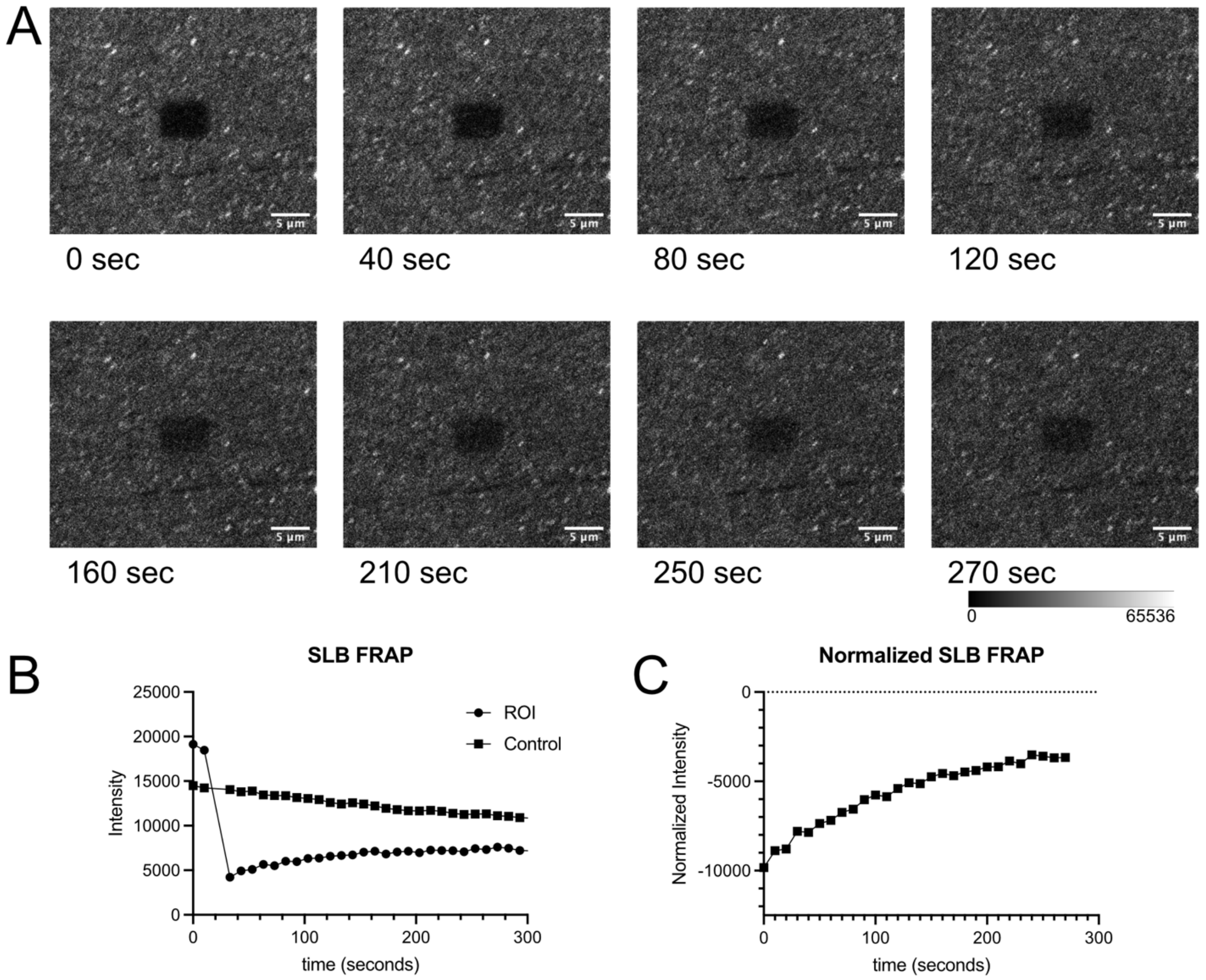


**Supplemental Figure 5: Supported lipid bilayers present mobile antigens.** (A) Representative image series acquired during FRAP experiment. (B) Raw fluorescence intensity from FRAP data. (C) Normalized fluorescence intensity presented in B.

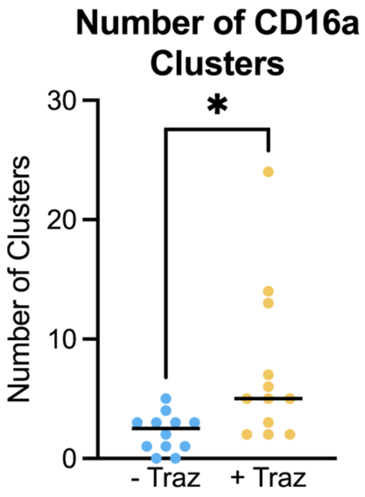


**Supplemental Figure 6: NK-92$^{CD16\text{-}SNAP}$ activate on SLB with Trastuzumab for increased number of clusters.** Comparison of the number of clusters identified by DBSCAN between NK cell synapses on SLBs without Trastuzumab (- Traz) and with Trastuzumab (+ Traz). MINFLUX microscopy data were collected across at least three independent experiments with a total of 12 cells analyzed. ** *p* = 0.0047.

### Materials and Methods

*Cell culture*

SKOV-3 cells (HTB-77) and NK-92 cells (CRL-2407), a malignant non-Hodgkin's lymphoma NK cell line, were purchased from ATCC (Manassas, VA, USA). SKOV-3 cells were maintained in Dulbecco's Modified Eagle's Medium (DMEM, Corning) supplemented with 10% (v/v) Fetal Bovine Serum (FBS, Gibco) and 1X Anti-Anti (Thermo Fisher). NK-92 cells were maintained in MyleoCult (Stemcell Technologies) supplemented with 10% FBS, 1X Anti-Anti, and 200 IU/mL IL-2 (Peprotech). Cells were maintained for less <20 passages and regularly tested for mycoplasma.

*Isolation of primary NK cells*

For assays that required primary NK cells, we utilized viably frozen PBMCs (a gift from Andrew Ward's lab). NK cells were isolated using an EasySep™ Human NK Cell Isolation Kit (StemCell Tech) according to the instructions provided in the kit. Freshly isolated NK cells were utilized immediately in ADCC assays (see below).

*CD16-SNAP plasmid*

The CD16-SNAP construct (Fig. S1) was synthesized and cloned into a transposon plasmid without reporter by Vector Builder.

*Cell line generation*

The NK-92$^{CD16\text{-}SNAP}$ cell line was constructed using the PiggyBac (PB) system. Approximately 3x $10^6$ NK-92 (ATCC) cells were co-nucleofected with a CD16-SNAP plasmid and the hyPBase plasmid carrying the transposase (VectorBuilder) at ratio of 3:1, respectively, using a Lonza P3 Primary Cell kit in approximately 100 µL medium and a Lonza 4D-nucleofector device with the program CA137. Post-nucleofection, cells were resuspended in complete medium and allowed to recover for 48 hours at 37˚C in 5% $CO_2$. Then half of the medium was removed and replaced with fresh complete medium. After 96 hours post-transfection, half of the medium was removed, and cells were resuspended again in a total of 10 mL of complete medium. After approximately 1-week post-transfection, cells were stained with an $\alpha$-CD16-AF488 (3G8 Biolegend, 1:20) antibody for 1 hour on ice and sorted using a Sony MA900 or Thermofisher Scientific Bigfoot cell sorter. Gating was performed on cells with fluorescence intensity greater than parental NK-92. After sorting, cells were placed in a 96 well plate U-bottomed plate in 200 µL of complete medium with IL-2 and expanded.

*Flow cytometry*

Confirmation of CD16a expression was performed by flow cytometry. Briefly, ~100,000 cells were placed in an Eppendorf tube and pelleted at 200 x g at 4°C for 5 minutes. Cells were then resuspended in 500 µL of PBS containing 1% (w/v) BSA. Cells were then washed twice with fresh, cold PBS with 1% BSA and then finally stained with an $\alpha$-CD16-AF488 antibody (3G8 Biolegend, 1:20) for 1 hour at 4°C. Cells were then washed twice with PBS with 1% BSA and resuspended in 200 µL for analysis on a CytoFLEX S (Beckman Coulter). Data analysis was performed using FlowJo.

*ADCC assay*

NK-92$^{CD16\text{-}SNAP}$ ADCC activity was measured using an Agilent xCELLigence Real-Time Cell Analysis (RTCA) SP impedance instrument that was setup in a stationary 37˚C incubator with 5% $CO_2$. On the first day of the experiment, SKOV-3 cells were detached using TrypLE™ Express medium, quenched with complete medium, and counted. 1 x $10^6$ cells were removed and pelleted at 250 x g for 10 minutes before being resuspended in complete Roswell Park Memorial Institute 1640 Medium (RPMI) at 1 x $10^5$ cells/mL. Meanwhile, 50 µL of complete RPMI was added to an E-Plate 96 PET plate (Agilent) and background impedance was measured. Then, 5,000 cells (50 μL) were added to each well and allowed to sit at room temperature for 30 minutes. The plate was then transferred to the instrument and impedance measured every 15 minutes for ~24 hours to allow cells to attach to the plate and establish baseline cellular impedance. The next day, 50 μL of Trastuzumab (Selleckchem) was prepared in complete RPMI medium and was added to yield a final concentration of 1 - 0.0016 μg/mL. While the antibody was allowed to bind, NK cells, NK-92, and/or NK-92$^{CD16-SNAP}$ cells were counted and pelleted at 150 x g for 10 minutes. Cells were then resuspended at 5 x $10^5$ cells/mL and 50 μL added to yield a final effector to target ratio of 5:1. The plate was then allowed to sit at room temperature for 30 minutes before being transferred into the instrument, and impedance was measured every 15 minutes for 24 hours. ADCC activity was determined using the Agilent xCELLigence software and curves were plotted using Prism (GraphPad).

*Sample preparation for confocal laser scanning microscopy*

Cover glasses (Corning, 18mm #1.5) were coated with 0.1% (w/v) PLL, α-CD18 antibody (5 μg/mL in PBS), or α-CD16/α-CD18 antibodies (5 μg/mL in PBS) for 30 minutes at room temperature. Cover glasses were then washed 3 times with PBS before plating NK-92$^{CD16-SNAP}$ cells (200 μL at ~1 x $10^6$ cells/mL) and placed in a humidified incubator at 37°C and 5% $CO_2$ for 30 minutes. Medium was then carefully removed by pipetting, and the sample was washed with PBS. Cells were then fixed with 4% PFA (w/v) for 15 minutes, permeabilized with 0.4% (w/v) Triton X-100 for 3 minutes and fixed again with 4% PFA for 15 minutes. Fixed cells were washed with PBS, quenched with $NH_4Cl$ (50 mM) for 5 minutes, and washed 3 times with PBS. Samples were then blocked with Image-IT (Invitrogen) for 30 minutes and washed 3 times with PBS.

Labeling of SNAP-tagged CD16a was performed with 1 μM AF647-SNAP (New England Biolabs) in PBS (0.5% w/v BSA, 1 mM DTT) for 50 minutes. Labeling solution was then removed, and samples were washed 2 times with PBS and allowed to wash in fresh PBS for 30 minutes.

PBS was then replaced with PBS containing 5% (w/v) BSA for ~2 hours. Samples were then incubated with an $\alpha$–pCD3$\zeta$ antibody (BD Bioscience, clone K25-407.69, AF488, 1:50) at 4°C overnight. Subsequently, samples were washed 3 times with PBS, post-fixed with 4% PFA for 3 minutes, fixative quenched with $NH_4Cl$ (50 mM) for 5 minutes, and finally washed 3 times with PBS before mounting the sample with Prolong Diamond (Invitrogen) for imaging.

*Liposome production*

Liposomes were prepared by dissolving 1-palmitoyl-2-oleoyl-glycero-3-phosphocholine (POPC) and (1,2-dioleoyl-sn-glycero-3-[(N-(5-amino-1-carboxypentyl)iminodiacetic acid)succinyl] (nickel salt)) DGS-NTA[$Ni^{2+}$] (Avanti) in chloroform at a 96:4 molar ratio. Lipids were dried under vacuum overnight, hydrated in PBS (20 mg/mL total lipid) with constant shaking for 2 hours at 37°C, then sonicated for 30 seconds. The resulting liposomes were extruded 14 times through 0.8 µm, 0.4 µm, 0.2 µm, and 0.1 µm filters using a Mini Extruder (Avanti) at room temperature.

*Supported lipid bilayer formation*

Supported lipid bilayers (SLB) were prepared on cover glasses (Deckgläser coverslips, #1.5). Cover glasses were first cleaned by sonication for 5 minutes in acetone, followed by 5 minutes of sonication in absolute ethanol, and a final sonication in MiliQ water. After each sonication step, the cover glasses were dried with a stream of nitrogen gas. After cleaning by sonication, the cover glasses were further cleaned by oxygen plasma in a Solarius plasma cleaner with a power of 20 W and a process pressure of 1.6 torr for 15 minutes before being attached to a 6 channel sticky-slide (Ibidi VI 0.4). After assembly of the channel slide, liposomes (50µL at 4 mg/mL) were deposited into each channel and incubated at room temperature for 20 minutes. The channels were then washed 10 times with PBS. SLBs were incubated with 100 µM $NiCl_2$ containing 1%(w/v) BSA for 20 minutes at room temperature. The samples were then washed 5 times with PBS and incubated with His-tagged HER2 (10 µg/mL; Acro Biosystems) and His-tagged ICAM-1 (1 µg/mL; Acro Biosystems) for 60 minutes at 37°C, after which the samples were washed 4 times with PBS.

*Immunological synapse formation on SLBs*

SLBs used for the preparation of NK cell immune synapses were prepared as described above, and subsequently incubated with either Trastuzumab (Selleckchem) at 10 µg/mL in PBS for 30 minutes at room temperature or PBS alone as a control, followed by three washes with PBS. Meanwhile, NK cells were counted, and 1 x $10^6$ cells were removed and pelleted at 100 x g for 10 minutes. Cells were then resuspended in serum-free RPMI and 100 µl were added to each channel of the prepared slide and incubated at 37°C in 5% $CO_2$ for 30 minutes. Medium was then removed, and samples were washed with PBS, fixed with 4%(w/v) PFA for 15 minutes, permeabilized with 0.4%(v/v) Triton X-100 for 3 minutes, and then fixed again with 4%(w/v) PFA for 15 minutes. Fixed cells were then washed with PBS, fixative was quenched with $NH_4Cl$ (50 mM) for 5 minutes, and the cells again washed 3 times with PBS. Samples were then blocked with Image-IT (Invitrogen) for 30 minutes and washed 3 times with PBS. Labeling of SNAP-tagged CD16a was performed with 2 µM AF647-SNAP (New England Biolab) in PBS containing 0.5%(w/v) BSA and 2 mM DTT for 50 minutes. Labeling solution was then removed, and samples were washed 2 times with PBS then allowed to wash in fresh PBS for 30 minutes. PBS was then replaced with PBS containing 5%(w/v) BSA for ~2 hours. Samples were next incubated with an $\alpha$-pCD3$\zeta$ antibody (BD Bioscience, clone K25-407.69, AF488) at 4°C overnight. Subsequently, samples were washed 3 times with PBS, post-fixed with 4%(w/v) PFA for 3 minutes, fixative was quenched with 50 mM $NH_4Cl$ for 5 minutes, the samples finally washed 3 times with PBS and fresh PBS was added before imaging.

*Preparing samples for fluorescence recovery after photobleaching (FRAP) sample preparation, collection and analysis*

SLBs were subjected to FRAP to determine bilayer fluidity. Samples were first prepared as described above. Subsequently, an α-Her2 antibody (R&D systems research grade trastuzumab biosimilar, AF488, 10μg/mL, 1:100) was added and incubated for 30 minutes at room temperature. Samples were then washed 3 times with PBS before proceeding to data collection. Fluorescently labeled SLBs were imaged on a Zeiss LSM 880 laser scanning confocal microscope using perfect focus with the following conditions: excitation at 488 nm (7% power) with a 493-630 nm emission detection window, 63x / 1.20 NA C-Apochromat water objective, 0.77 μs pixel dwell time, 800V gain offset, and 0.099 μm pixel size. Images were acquired every 30 seconds for 30 scans. After two scans, a region of interest (ROI) of ~25 $\mu m^2$ was photobleached (100% power, 10 μs pixel dwell time, for a single scan) and subsequent fluorescence recovery was recorded.

*Confocal image collection*

Confocal images of NK cells stained for SNAP-CD16 and pCD3ζ were acquired on a Zeiss LSM 780 laser scanning confocal microscope under the following conditions: excitation at 488nm and 633nm (10% power) emission with detection windows of 493-628 nm and 638-755 nm, respectively, a 63X / 1.40NA plan-apochromat oil objective with a pixel dwell time of 4.08 μs, line averaging of 2, pixel size of 0.09 μm, and a master gain of 800V. Z-stacks of ~2 μm were collected with a step size of 200 nm for each cell to cover the height of the immunological synapse. The in focus z-plane was then used for quantification.

*MINFLUX sample preparation*

Samples for MINFLUX data collection were stained for SNAP-CD16 as described above for confocal sample preparation. After removal of excess SNAP-AF647 staining solution, cells were labeled with phalloidin (Invitrogen, AF488, 1:200) for 60 minutes and washed 3 times with PBS. Before mounting, 150 nm diameter gold fiducials (AUROlite™ Au/$TiO_2$, STREM) were added to the samples for 5 minutes and then washed to remove unbound beads away 3 times with PBS.

The antibody coated cover glasses were placed onto glass slides featuring a cavity well filled with imaging buffer (50 mM Tris-HCl, 10 mM NaCl, 10% (w/v) glucose, 10 mM cysteamine, 40 µg/mL bovine-liver catalase and 100 µg/mL glucose oxidase from *Aspergillus niger*, type VII, pH 8.0 (GLOX buffer)) and pressed down to remove excess buffer. The cover glasses were sealed onto the slide using Elite Double 22 dental epoxy (Zhermack) and mounted onto the MINFLUX microscope. For samples on SLBs, PBS in the channels was replaced with GLOX buffer before sealing the channels with parafilm and mounting the slides on the MINFLUX microscope.

*Image collection MINFLUX*

All MINFLUX data were acquired on a commercial MINFLUX 3D microscope equipped with a 640 nm continuous-wave (CW) laser for excitation and a 405 nm CW laser for activation, using the Imspector software with MINFLUX drivers (Abberior Instruments). A field of view with three or more gold fiducials was chosen to allow for the active sample stabilization unit to lock onto the chosen spatial set point with live feedback correction based on the near-infrared scattering signal from gold fiducials. Throughout all MINFLUX recordings, the mean square displacement of the sample position, relative to the stabilization set point was less than 3 nm along all three axes.[19] A 3 x 3 µm ROI at the bottom of the cell was chosen for MINFLUX imaging. The majority of the on-state fluorophores within the ROI were driven into the dark state through iterative confocal scanning of the 640nm laser (8 µW power, 0.02 µm pixel size, 10 µs pixel dwell time). The same ROI was then imaged for 3 hours using the company's standard 2D MINFLUX imaging sequence as provided by the manufacturer (hexagonal targeted coordinate pattern (TCP) with a localization range (L) reaching 40 nm in the final iteration) with a 640 nm excitation power of 13 µW set in the software interface. The 405 nm laser was manually increased incrementally throughout the course of the experiment to a max of 320 nW. During the last iteration of the MINFLUX targeting sequence, the laser power is ramped up to by a factor of six.

MINFLUX data reconstruction was performed using a custom written python scrip. Raw data were reconstructed with 10 nm circles. Reconstructions containing trace centers were reconstructed with 5 nm circles.

*Data analysis confocal*

All confocal images were analyzed using Imaris 10.1 particle extraction with dynamic background. Identified particles were then filtered to remove small particles of a few pixels (manual threshold 337 and 1908 for CD16a and pCD3ζ, respectively). Filtered particles were counted, and the area and mean fluorescence intensity were averaged to provide a single data point per cell.

*Data analysis MINFLUX*

MINFLUX data were analyzed using a custom python script. Briefly, data were filtered to remove individual localizations with effective frequencies in offset (efo, background-corrected emission rates) greater than 50,000 Hz and center frequency ratios (cfr, ratio between photon count detected during TCP center exposure and photon count detected during TCP offset exposures) less than 0.95, and traces (groups of individual localizations originating from the same emission event) with fewer than 3 localizations were removed to remove traces containing multiple fluorophores and low confidence traces. The center of each trace was then found using DBSCAN[44] with a search radius of 4 nm and a threshold of 3 localizations. The 2D-coordinates for each center were then used for further analysis. Nearest neighbor analysis was performed using the ball tree algorithm in Sci-kit learn.[46] The nearest neighbor of isolated pairs was determined by identifying clusters of 2 trace centers in a 40 nm search radius using DBSCAN and filtering out clusters with more than 2 trace centers. The nearest neighbor of the filtered data was then determined using the ball tree algorithm in Sci-kit learn.[46] Ripley H analysis was completed as previously described, implemented as a custom function in python.[37]

*Statistics*

Confocal data were collected over at least three independent experiments with >30 cells per experiment. Statistical significance was calculated using either one-way ANOVA Kruskal-Wallis test comparing all samples or Mann Whitney test. Calculations were carried out using PRISM 10. MINFLUX microscopy data were collected across at least three independent experiments with a total of 12 cells analyzed. Statistical significance was calculated using the Mann Whitney test to compare number of clusters.
